# Single-shot Volumetric Chemical Imaging by Mid-Infrared Photothermal Fourier Light Field Microscopy

**DOI:** 10.1101/2022.06.24.497516

**Authors:** Yi Zhang, Danchen Jia, Qianwan Yang, Yujia Xue, Yuying Tan, Zhongyue Guo, Meng Zhang, Lei Tian, Ji-Xin Cheng

**Affiliations:** Department of Physics, Boston University, MA, 02215, Boston; Department of Electrical and Computer Engineering, Boston University, MA, 022153, Boston; Department of Biomedical Engineering, Boston University, MA, 022154, Boston; Photonics Center, Boston University, MA, 02215, Boston

**Author notes:** equal contributions (Yi Zhang, Danchen Jia, Qianwan Yang).

## Abstract

Three-dimensional molecular imaging of living organisms and cells plays a significant role in modern biology. Yet, current volumetric imaging modalities are largely fluorescence-based and thus lack chemical content information. Mid-infrared photothermal microscopy as a new chemical imaging technology provides infrared spectroscopic information at sub-micrometer spatial resolution. Here, by harnessing thermosensitive fluorescent dyes to sense the mid-infrared photothermal effect, we demonstrate mid-infrared photothermal Fourier light field (MIP-FLF) microscopy for single-shot volumetric infrared spectroscopic imaging at the speed of 8 volumes per second and sub-micron spatial resolution. Protein contents in bacteria and lipid droplets in living pancreatic cancer cells are visualized. Altered lipid metabolism in drug-resistant pancreatic cancer cells is observed with the MIP-FLF microscope.

## 1. Introduction

Volumetric imaging plays a prominent role in life science due to its ability of visualizing 3D architecture of biological systems ranging from whole-brain to subcellular level^1-3^. Volumetric fluorescence imaging is routinely performed by optical sectioning and z-axis scanning with either confocal microscopy^4^ or two-photon laser scanning microscopy^5^. However, these scanning-based approaches are time-consuming and require a precise mechanical 3D positioning device. In cases where photobleaching is a concern, repeated sectioning only exacerbates this problem by increasing exposure. Fast focus scanning methods using electrically tunable lens^6^ or spatial light modulator^7^ mitigate the issue by improving scanning speed, but hinder the imaging contrast due to a drawback of out-of-focus background. Light-sheet fluorescence microscopy physically eliminates the background by illuminating the sample only with a thin sheet of light from the side of the specimen for optical sectioning^8-10^. Also, the excitation is confined to the focal plane, thus boosting detection efficiency while reducing photobleaching and photodamage. Various scanning-based strategies discussed above promote the imaging speed of volumetric fluorescence imaging^11^, but still cannot achieve video-rate due to the requirement of mechanical movement.

To further capture the fast-moving organelles or study dynamic activities in living cells, single-snapshot light field microscopy emerged as a scanning-free, scalable method that allows for high-speed, volumetric imaging^12^. Specifically, a microlens array is used to capture 3D structure of objects in a single snapshot. Recently developed Fourier light-field (FLF) microscopy achieved high-quality volumetric imaging by recording the light field in the Fourier domain, which allows jointly allocating the spatial and angular information of the incident light in a consistently non-overlapping manner, effectively avoiding any artifacts in conventional light field systems^13-16^. These advances in both imaging capability and computation speed promise further development of FLF microscopy toward high-resolution, volumetric data acquisition, analysis and observation at the video rate. However, these methods are fluorescence based, lacking the chemical content information about the subjects.

Vibrational spectroscopic imaging based on molecular fingerprints is able to offer chemical content and molecular structural information about an organism in a label-free manner^17^. Along this direction, coherent Raman scattering microscopy has been developed and has found broad applications in revealing recording neural activities^18^, cancer metastasis^19^, brain metabolic activity^20^, and so on. Bessel beam based Stimulated Raman projection tomography has been developed to quantify the total chemical composition of a 3D object^21^. More recently, volumetric chemical imaging on the nanoscale is enabled by the integration of stimulated Raman scattering microscopy and expansion microscopy^22, 23^. Yet, the detection sensitivity of Raman based vibrational microscopy is ultimately limited by its small cross section.

The infrared absorption offers a cross section that is eight orders of magnitude larger than Raman scattering. Yet, infrared microscopy lacks depth resolution, which prohibits accurate decomposition of cellular dry mass density into independent biomolecular components. Recently developed mid-infrared photothermal (MIP) microscopy exceeds the diffraction limit of infrared microscopy and allows three-dimensional chemical imaging in a confocal scanning manner^24^. To improve the 2D imaging speed and achieve large scale imaging, wide field MIP microscopy is demonstrated via a virtual lock-in camera approach^25^. By measuring the mid-infrared photothermal effect in a quantitative phase microscope, both 2D and 3D bond-selective phase imaging has been demonstrated^26-29^. Yet, the speed (∼50s per volume) is insufficient to capture chemical information in a highly dynamic living system.

Here, we present a mid-infrared photothermal Fourier light-field (MIP-FLF) microscope that allows single-shot chemical imaging of living cells. We harness thermosensitive fluorescent dyes to sense the mid-infrared photothermal effec^t30, 31.^ The fluorescence intensity can be modulated at the level of 1% per Kelvin by mid-IR absorption, which is 100 times larger than the thermal modulation of scattering intensity. Moreover, the fluorophores can target specific organelles or biomolecules, thus augmenting the specificity of photothermal imaging. Importantly, this method is fully compatible with the FLF fluorescence imaging. By recording photothermal modulation of fluorescence emission in an FLF microscope, we achieved an infrared spectroscopic imaging rate of 8 volumes per second, at a lateral resolution of 0.5 to 0.9 µm and an axial resolution of 0.8 to 1.1 µm. This speed is faster than reported confocal MIP microscopy^24^ or MIP-based optical diffraction tomography^28^ by two orders of magnitude. With these advancements, we demonstrate single-shot volumetric bond-selective imaging in living cells, where the lipid content is used as a marker to determine the drug resistance of cancer cells.

## 2. Methods

### 2.1. MIP-FLF Microscope

The MIP-FLF microscope is a pump and probe imaging system (**Fig. 1a**). A pulsed visible probe beam passes through a FLF microscope to capture fluorescently labeled sample, while the IR absorption information is coded by a nanosecond pulsed mid-IR laser. The raw MIP-FLF images are acquired by synchronizing the IR pump pulses, the visible probe pulses, and the camera exposure (**Fig. 1b**). After acquiring each pair of “IR-on” and “IR-off” images, a 3D deconvolution is performed that incorporates the point spread function (PSF) to generate the final 3D chemical image (**Fig. 1c**).

**Figure 1.**
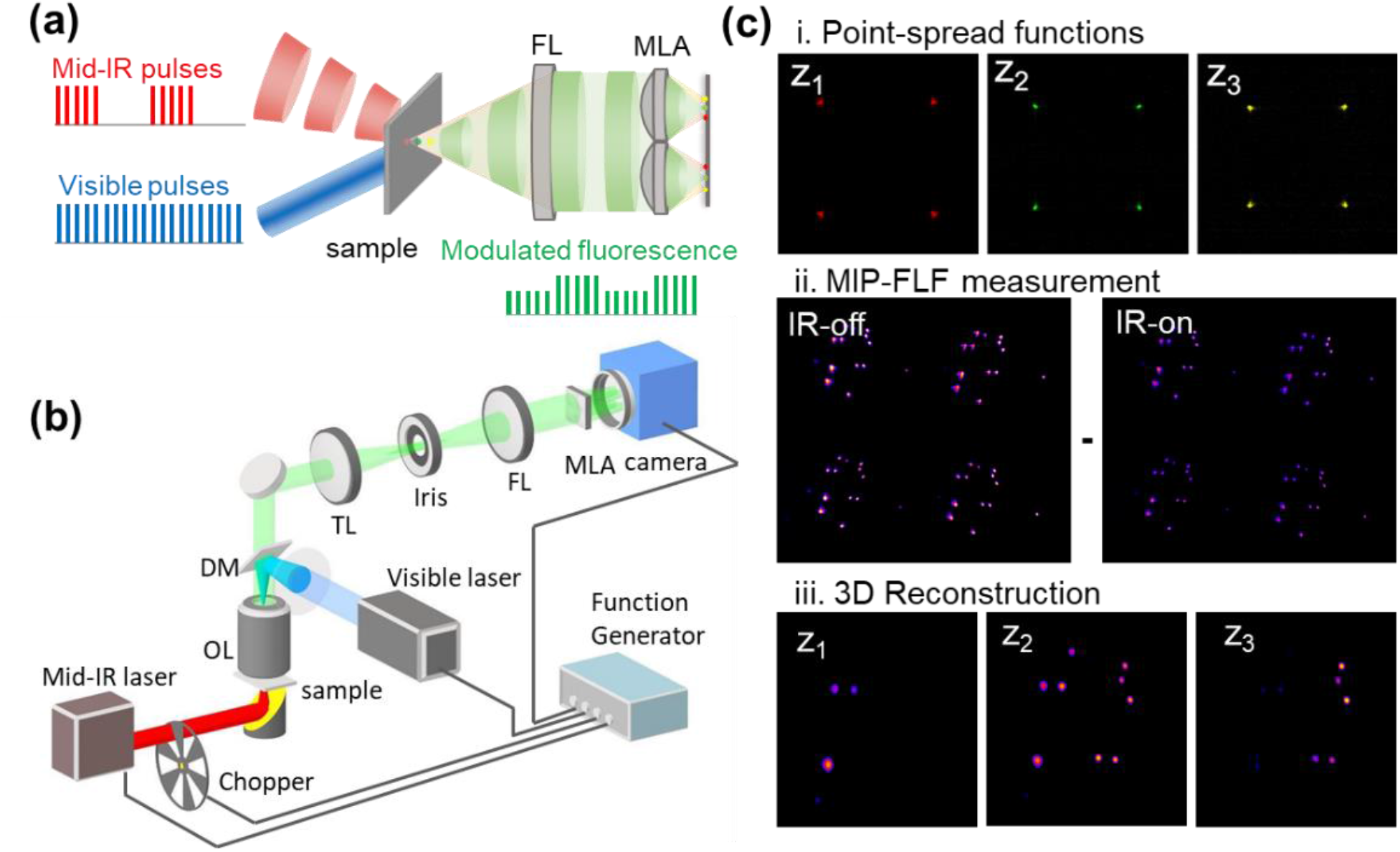
MIP-FLF imaging system and procedures. (a) Principle of MIP-FLF microscope. The modulated fluorescence emission carries IR information coded by mid-IR pulses. Each single snapshot image contains 3D information introduced by a microlens array (MLA) placed at the Fourier plane. FL, Fourier lens. (b) Experimental setup. The pump beam is a nanosecond mid-IR laser, modulated by an optical chopper and weakly focused on the sample by a gold-coated parabolic mirror. The probe beam is a 520 nm nanosecond laser and sent through the MLA placed at the back focal plane of a FL. The tube lens and the FL formed a 4f system. OL, objective lens. DM, dichroic mirror. (c) Procedures of MIP-FLF measurements. (i) Example measured PSFs of the system. (ii) Raw MIP-FLF measurement contains a pair of IR-on and IR-off images. (iii) 3D MIP-FLF reconstruction by deconvolution between 2D raw MIP-FLF image and the measured PSFs. The samples were Rhodamine 6G stained *Staphylococcus Aureus* (*S. aureus*) bacteria prepared on a tilted silicon substrate.

The schematic of the MIP-FLF microscope is shown in **Fig. 1b**. The infrared pulses are generated by a tunable optical parametric oscillator laser (Firefly-LW, M Squared Lasers), ranging from 1175 to 1800 cm^-1^, operating at 20 kHz repetition rate and 50 ns pulse duration. An optical chopper (MC2000B, Thorlabs) modulates the IR pulses to accommodate the acquisition speed of the camera. A gold-coated off-axis parabolic mirror with a focal length of 25.4 mm weakly focuses the pump beam at the sample.

On the FLF microscope, a 2×2 microlens array is used to image the 3D fluorescence as 2×2 projection views on the camera plane. The FLF imaging is implemented on an epifluorescence microscope using a 100×, 0.95NA objective lens (PLFLN100X; PLAN FLUOR 100X DRY OBJ, NA 0.95, WD 0.2). The samples are excited with a 520-nm nanosecond laser (NPL52C, Thorlabs) at 20 kHz repetition rate. The generated fluorescence emission is collected with a dichroic mirror (DMLP550T, Thorlabs), a 550-nm long-pass filter (FEL0550, Thorlabs) and a tube lens (TTL180-A, Thorlabs). The field of view (FOV) is adjusted by an iris placed at the native image plane to avoid overlapping light field signals on the camera plane. A Fourier lens (FL, *f*_*FL*_= 150 mm, Thorlabs) performs optical Fourier transform of the image at the native image plane, forming a 4f system with the tube lens (TL). A microlens array (MLA, APO-Q-P3000-R23.5, advanced microoptic systems GmbH) is placed at the back focal plane of the FL, thus conjugated to the back pupil of the objective. The raw FLF images are recorded on a CMOS camera (CS235MU, Thorlabs) at the back focal plane of the MLA.

Before recording MIP-FLF images, a one-time calibration of the system 3D PSF is required. To perform the calibration, we first prepared a point source phantom with resolution limited fluorescence beads (P7220, 175 nm diameter, excitation/emission 540/560 nm, PS-Speck™). We adjusted the concentration of the beads to ensure only one bead is in the imaging area. The PSFs were recorded by scanning the point source along the axial direction with a 100 nm step size (**Fig. 1c-i**).

To record the MIP-FLF images, a pulse generator (Emerald 9254, Quantum Composers) triggers the optical chopper, the 520 nm nanosecond laser and the CMOS camera with a master clock signal of 20 kHz from the mid-IR laser. The camera sequentially measured the “IR-on” and “IR-off” FLF images (**Fig. 1c-ii**). MIP-FLF signals were extracted by subtracting the two images. The 3D objects were then reconstructed through a deconvolution algorithm using the 2D MIP-FLF measurement and the calibrated PSFs. 3D chemical information was attained by scanning the wavenumber of mid-IR laser during MIP-FLF measurements or concentrating on the vibrational frequencies of certain chemical bonds.

### 2.2. FLF reconstruction algorithm

The forward model of the FLF system can be written as the following form^13^

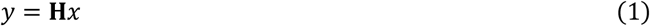

where y denotes the 2D sensor measurement, x is the 3D object, **H** is the forward convolution matrix determined by the experimentally calibrated 3D PSF. The 2D measurement is treated as the axial sum of the 2D convolution between the object “slice” at each depth and the corresponding depth-dependent PSF.

The reconstruction algorithm solves a regularized least squares optimization

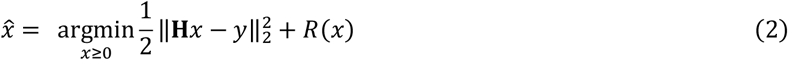

Here, R refers to regularization terms that includes *l*_1_-norm and 3D total variation to promote sparsity of the 3D solution. We solve the optimization by adopting the algorithm based on the alternating direction method of multipliers (ADMM)^16^.

### 2.3. Bacteria, cancer cell culture, and lipid droplets staining

*S. aureus* was stained with 10^−4^ mol/L Rhodamine 6G (R6G) for 20 minutes and then dried on a tilted silicon substrate. Pancreatic cancer MIA Paca-2 cells were purchased from the American Type Culture Collection. The cells were cultured in Dulbecco’s Modified Eagle Medium (DMEM) medium supplemented with 10% Fetal Bovine Serum and 1% Penicillin-Streptomycin. All cells were maintained at 37 °C in a humidified incubator with 5 % CO_2_ supply. For lipid droplets staining, cells were cultured within 3 μmol/L lipi-red (Dojindo Lipi-Series) dissolved in serum-free medium at 37 °C for 30 minutes, following three times of Phosphate-Buffered Saline (PBS) wash. After staining, cells were washed with deuterated PBS solution and sandwiched the cells between glass coverslip and silicon substrate for MIP-FLF imaging. In the experiments of ^13^C fatty acid treatment, MIA Paca-2 cells were cultured with 25 g/L ^13^C isotopic labeled fatty acids mixture for 24 hours.

## 3. Results

### 3.1. System characterization

To characterize the spatial resolution of the MIP-FLF system, we prepared an agarose film filled with fluorescent beads of 175 nm diameter. The fluorescence beads were distributed in 1% agarose gel through sonication. The gel then formed a thin film on a silicon substrate. The raw FLF image contains four (2×2) elemental images (**Fig. 2a**), which are the perspective views containing different spatial and angular information captured by the MLA. Based on the forward model, the 3D object (**Fig. 2b**) was reconstructed through our deconvolution algorithm (**Fig. S1**, Methods). The full width at half-maximum (FWHM) of the reconstructed beads at varying depths was measured using Gaussian fitting (see **Fig. S2**), resulting in a lateral resolution of 0.5-0.9 μm and an axial resolution of 0.8-1.1 μm. The measured image volume was ∼60 μm × 60 μm × 4 μm due to the trade-off between spatial resolution and FOV in FLF microscopy. The axial resolution was determined by the lateral translation of the objects from varying depths on the camera plane, which was demonstrated in the depth-resolved PSFs of the system. By comparing the MIP-FLF reconstruction results with widefield epifluorescence images at varying depths (**Fig. 2d**), we confirmed the imaging fidelity of single-shot 3D MIP-FLF microscopy, which is capable of imaging with submicron spatial resolution. Furthermore, the MIP-FLF microscope provided significant improvement in the axial sectioning capability compared with widefield fluorescence (**Fig. 2d**).

**Figure 2.**
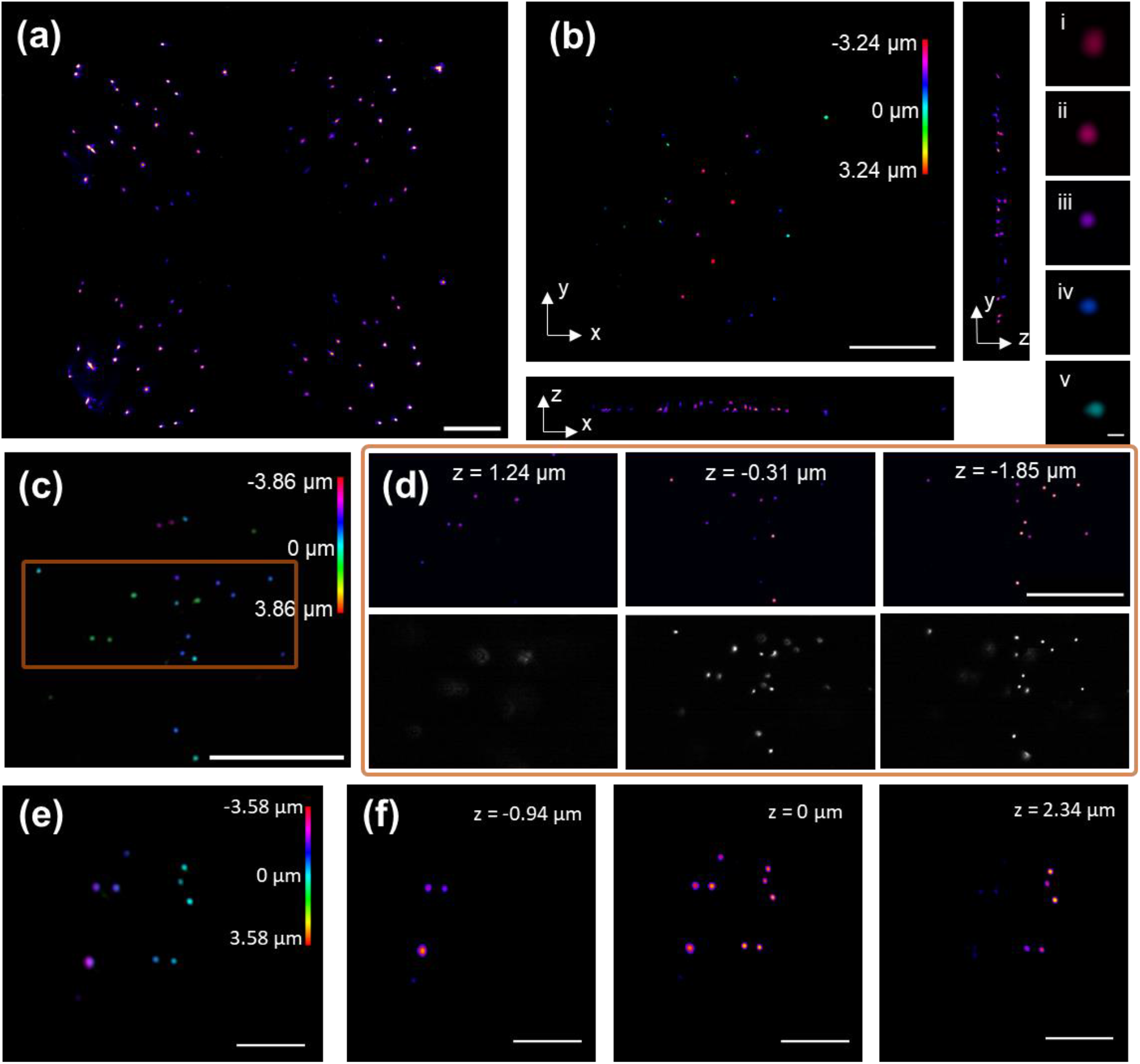
Characterization of MIP-FLF system with 175 nm fluorescent beads and S. aureus. (a) Raw FLF measurement, and (b) 3D reconstructed image of 175 nm fluorescent beads distributed in 3D agarose gel, and its x-z and y-z projection. The inset images (i-v) are zoomed-in images of (b) at z = −1.8 to 1.0 µm, with FWHM of 0.5 to 0.9 µm and 0.8 to 1.1 µm in the lateral and axial direction, respectively (see Fig. S2). (c) 3D reconstructed fluorescent beads and (d) zoomed-in reconstructed slices at varying depths and corresponding widefield images with the same FOV. (e) 3D reconstructed FLF image and (f) reconstructed MIP images of varying depth at 1650 cm-1 of R6G stained S. aureus prepared on a tilted Si substrate. Scale bar, 20 µm; scale bar of (b) insets, 1 µm.

We further demonstrated MIP-FLF imaging of biological samples using S. aureus to confirm the spectral fidelity of the system. In the MIP measurement, fluorescence intensity was modulated by mid-IR pump beam due to the photothermal effect (**Fig. 1c-ii**). MIP signals were thus extracted through subtraction of sequentially acquired IR-on and IR-off frames of the FLF image stack. The MIP-FLF reconstructed 3D image stack showed the mid-IR absorption mapping of S. aureus at the Amide-I band (1650 cm-1) in **Fig. 2e**. Notably, the volumetric MIP-FLF image stack was reconstructed from a single-shot FLF image, including one IR-on and one IR-off frame. Consequently, the MIP-FLF system is capable of video-rate 3D chemical imaging. As a result, MIP-FLF microscopy is able to capture fast-moving components and study activities in dynamic living cells with chemical specificity, high throughput, as well as high spatial resolution.

### 3.2. 3D chemical imaging of lipid droplets in cancer cells

To demonstrate the capability of FLF-MIP system for live cell imaging, we imaged MIA-Paca2 cells stained with lipi-red (see Methods). The bright spots in **Fig. 3b** represent lipid droplets in a single MIA-Paca2 cell. By using a 1744-cm-1 mid-IR pump beam to excite the C=O bonds in esterified lipids, MIP modulation of lipid droplet fluorescence was demonstrated by the comparison between IR-on and IR-off frames (**Fig. 3a**, i and ii, respectively). After reconstruction, 3D distribution of lipid droplets in a single cell was mapped in **Fig. 3b** with bond-selectivity, covering a volume of ∼60 μm×60 μm×4 μm. A few reconstructed depth slices are shown in **Fig. 3c**. Here, the depth-resolved MIP-FLF images of lipi-red stained MIA-Paca2 showed rich lipid contents in cancer cells, except for the nuclei area. The 3D reconstructed MIP-FLF images with submicron level axial resolution and video acquisition rate show greater potential in quantitative chemical analysis than previous MIP microscopy in 2D manner. Furthermore, using this ultrafast 3D imaging method in living cells with chemical specificity, we are capable of studying lipid metabolism in cancer cells, such as fatty acid uptake that is beyond the reach of FLF microscopy.

**Figure 3.**
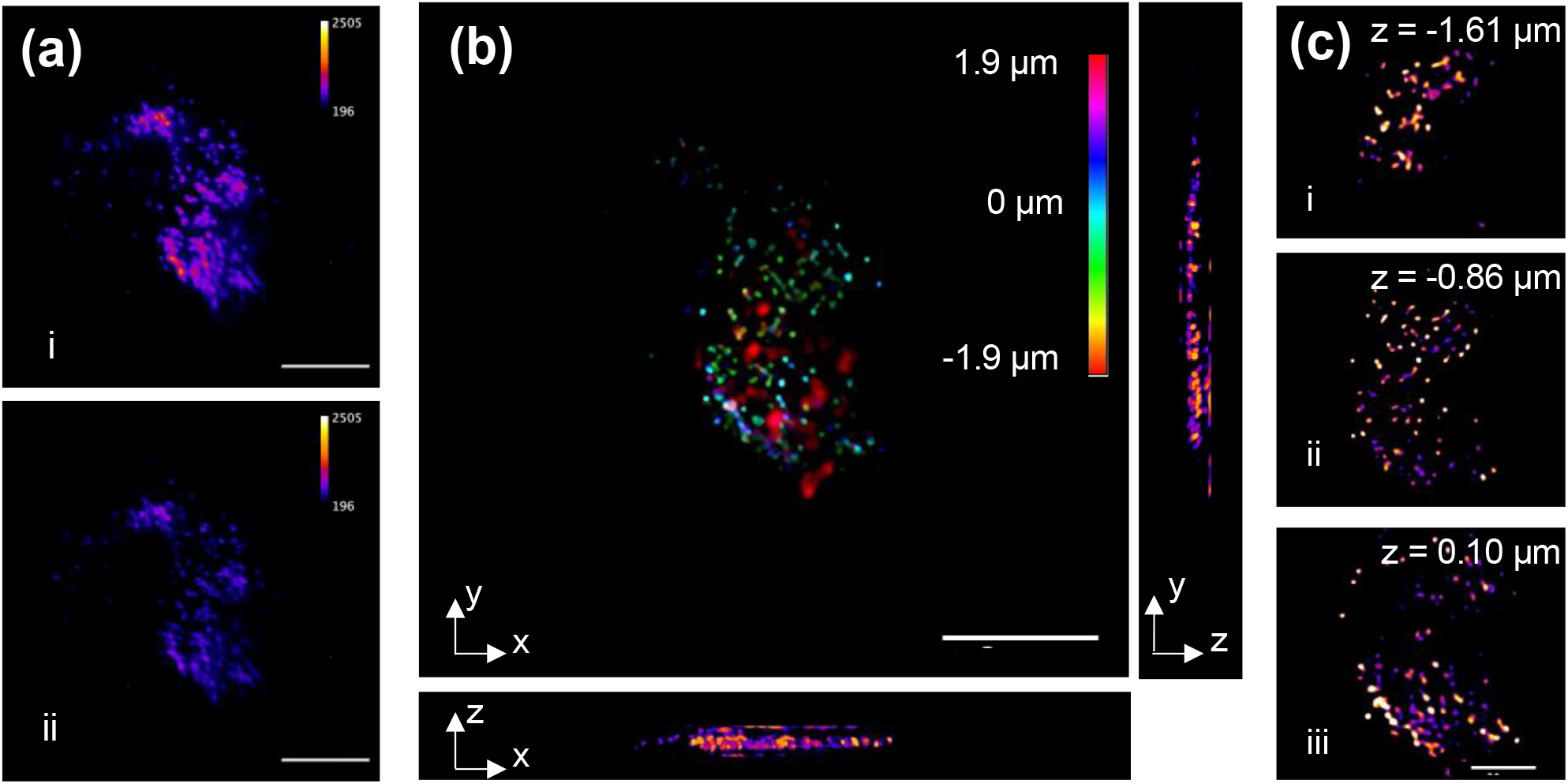
MIP-FLF imaging of lipid droplets in a MIA Paca-2 cell. The IR beam was tuned to 1744 cm-1 for the excitation of C=O bonds. (a) Raw FLF elemental images at (i) IR-on and (ii) IR-off state. (b) 3D reconstructed MIP image and its x-z and y-z projections. (c) 3D reconstructed MIP images at varying depth. Scale bar of (a, b), 20 µm; scale bar of (c), 10 µm.

### 3.3. MLP-FLF imaging of lipid metabolism in drug-resistant cancer cells

Lipids are essential building blocks synthesized by complex molecular pathways, from glucose or fatty acid uptake, and are stored as lipid droplets in cells. Currently, ^13^C metabolic flux analysis is the preferred tool for quantitative characterization of metabolic phenotypes in mammalian cells^32,33^. Cells can take up long chain free fatty acids (FFA) *in vivo* from the non-protein bound ligand pool in extracellular fluid^34^. To quantitatively study fatty acid uptake in cancer cells, we treated MIA Paca-2 cells with ^13^C isotopically labeled fatty acids. As seen in **Fig. S3**, Fourier-transform infrared spectroscopy (FTIR) of ^13^C labeled fatty acid mixture showed a ∼30 cm^-1^ peak shift to lower wavenumber compared with ^12^C palmitic acid (major contents in the ^13^C fatty acid mixture). Incubating MIA Paca-2 cells with ^13^C isotopic labeled free fatty acid results in the formation of intracellular neutral lipid. Consequently, the MIP peak of lipid droplets formed from ^13^C fatty acid treatment is expected to display a ∼30 cm^-1^ peak shift from that for endogenous, de novo synthesized lipids (1744 cm^-1^). As seen in **Fig. 4c**, the MIP spectra (solid curves in **Fig. 4c**) were generated by Lorentzian fitting from MIP signals of MIA Paca-2 cells (N=3) heated by mid-IR pulses ranging from 1690 to 1745 cm^-1^. Since the sampling rate in frequency domain of MIP-FLF imaging were limited by the photobleaching effect of the fluorescent dye, MIP-FLF images were collected only at six mid-IR wavenumbers for each cell. Lipid droplets in ^13^C fatty acid treated cells (red) showed MIP peaks shift to shorter vibrational frequency, while those without ^13^C fatty acid treatment (green) remained at 1744 cm^-1^. Besides, cells treated with ^13^C fatty acid of high concentration (red) displayed higher MIP signals at ^13^C=O peak than those of low concentration (light red). This indicates that our isotopically labeled MIP-FLF imaging method is capable of quantitative study of lipid metabolism. Taking the ratio of the MIP intensity between 1704 cm^-1^ and 1744 cm^-1^, the lipid chemical components of MIA-Paca2 were demonstrated in **Fig. 4a, b**, where the red colormap represented lipid droplets formed from fatty acid uptake and the green color mapped endogenous lipid. Here, the ratio colormap of MIA Paca-2 confirmed rich lipid contents in cancer cells from fatty acid pathway, while the restored MIP spectra demonstrated the fatty acid uptake quantitatively.

**Figure 4.**
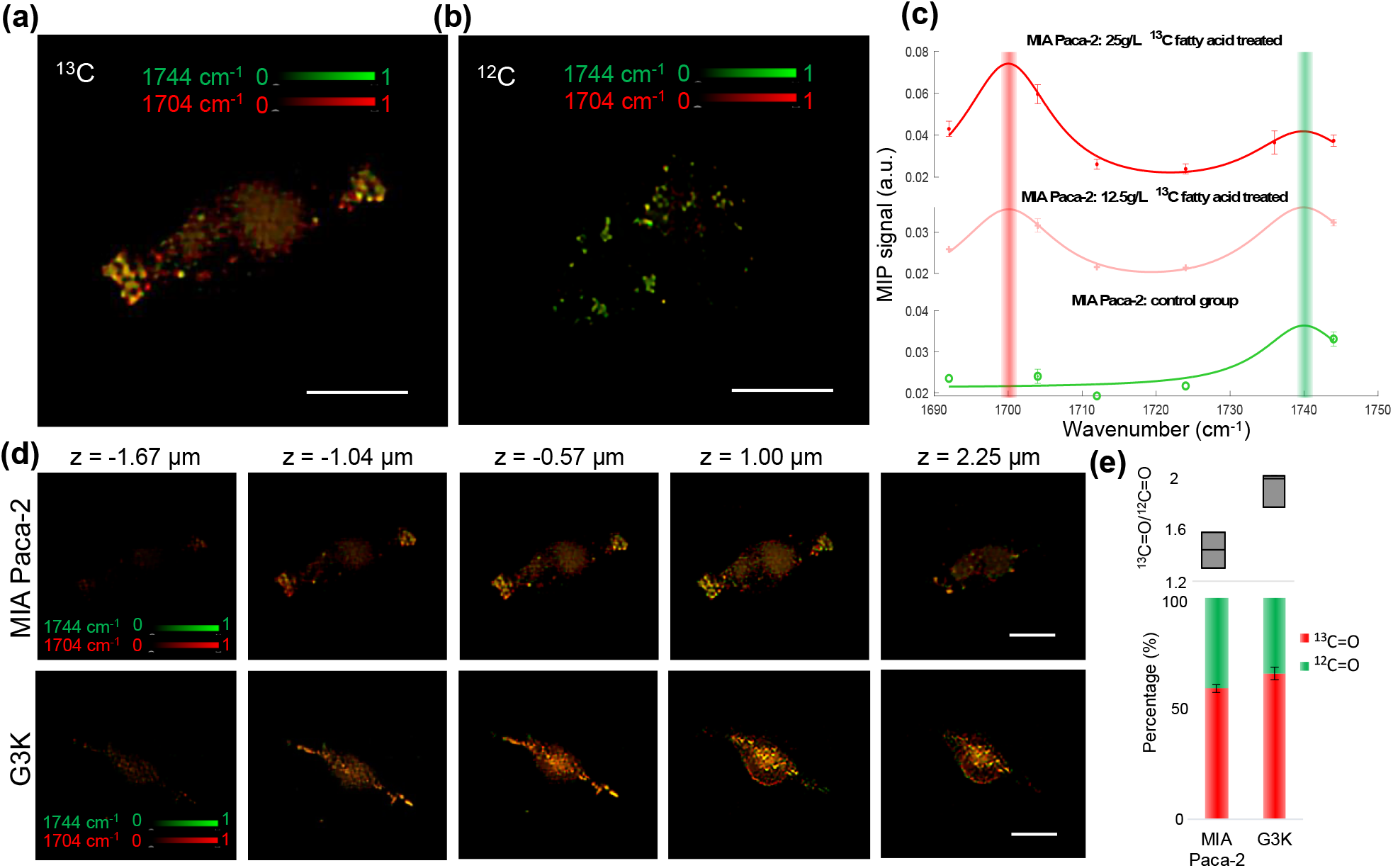
MIP-FLF imaging of lipid droplets in MIA Paca-2 and G3k cells treated with 13C fatty acids. (a-b) MIP intensity from 3D MIP-FLF reconstructed stack of 13C fatty acid treated MIA-Paca-2 cells (a) and the control group (b) without fatty acid treatment imaged at 1704 cm-1 (red) overlayed with that from 1744 cm-1 (green). (c) MIP spectra of MIA Paca-2 cells treated with (red, 25g/L, light red, 12.5g/L) and without (green) 13C fatty acid. The solid curves are Lorentzian fitted MIP spectra (N=3). (d) MIP intensity colormap of 13C fatty acid treated MIA-Paca-2 cells (top) and G3K cells (bottom) (green, 1744 cm-1, red, 1704 cm-1) at varying depths. (e) The ratio of MIP intensity between 13C=O at 1704 cm-1 and 12C=O at 1744 cm-1 (top). Statistical analysis of lipid content in 13C fatty acid treated MIA Paca-2 cells (N=3) and G3K cells (N=3). Scale bar 20 µm.

Next, we used the MIP-FLF ratio image to study altered fatty acid metabolism in drug-resistant pancreatic cancer cells by comparing the gemcitabine-sensitive MIA PaCa-2 cells and gemcitabine-resistant G3K cells (See Methods). Based on the lipid pathway tracing method discussed above, we can distinguish between exogenous and endogenous lipid of living cells. Here, both G3K group and MIA Paca-2 group were treated with ^13^C fatty acids with the same concentration and culture time. In the depth-resolved MIP-FLF reconstructed image stacks (**Fig. 4d**), drug-resistant G3K group shows higher MIP intensity at 1704 cm^-1^ (red) than the MIA Paca-2 cell. As compared in the statistical analysis of the lipid droplets in MIA Paca-2 (N=3) and G3K (N=3) cells (**Fig. 4e, S4**), G3K group displayed exceedingly more ^13^C=O than MIA Paca-2 group. As a result, we concluded that gemcitabine-resistant G3K cells more rely on exogenous fatty acids than gemcitabine-sensitive MIA PaCa-2 cells. Meanwhile, the G3K cells showed distinctly longer protrusions than MIA Paca-2, and the protrusions were rich in lipid droplets. These observations can also serve as markers to determine drug resistance of cancer cells.

## 4. Discussion

We developed MIP-FLF microscopy that allows single-shot volumetric chemical imaging with infrared spectroscopic information at an 8-Hz volume rate. The image volume was 60 μm×60 μm×4 μm with a lateral resolution of 0.5-0.9 µm and an axial resolution of 0.8-1.1 µm. FLF microscopy provides an unprecedented 3D imaging speed that cannot be achieved with the confocal microscope, which takes a few seconds to scan the sample voxel by voxel. The MIP sets additional requirement for the confocal scanning when detecting the fluorescence intensity modulation induced by the mid-IR pulse. The pump mid-IR laser can only operate at a limited repetition rate, which is determined by not only the laser pulse repetition rate, but also the thermal decay time to avoid heat accumulation. In addition, to get enough SNR, long pixel dwell time is used experimentally and thus further decreases the scanning speed and makes confocal measurement not an optimal method for high-speed volumetric chemical imaging. The light sheet microscopy has several benefits over the confocal laser scanning microscopy. The plane-scanning speed is faster than point-scanning speed and also diminish the photobleaching issue. However, all these scanning-based methods are not capable of imaging 3D volume in a single snapshot.

The MIP-FLF system is compact and simple, since it does not require any mechanical movements. Meanwhile, the single-shot volumetric system is more stable than scanning-based methods, which plays a significant role in MIP detections. During the MIP measurements, mechanical fluctuation induced by scanning or rotation will introduce strong artifacts after the subtraction of images at IR-on and IR-off states, thus is detrimental to the restoration of spectroscopic information. Furthermore, the ultrahigh speed of MIP-FLF imaging also mitigate the laser fluctuation issue during MIP detections.

In the reported MIP-FLF microscope, single-shot chemical 3D volumes captured in half of the camera frame rate contains volumetric chemical information in video rate. However, real-time volumetric spectroscopic imaging is still challenging due to the long reconstruction time. Recently, deep learning framework has emerged as the state-of-the-art to perform fast and reliable reconstruction for various volumetric fluorescence microscopes^35-37^. We anticipate that our system could achieve video-rate reconstruction speed by replacing the current iterative method with deep learning framework, which is especially beneficial for biomedical application.

In summary, via the development of MIP-FLF microscopy, we demonstrated single-shot volumetric chemical imaging of lipid contents in living cells and further studied the lipid metabolism in drug-resistant cancer cells with infrared spectroscopic selectivity. Specifically, we distinguished lipids from exogeneous fatty acids versus those from de novo synthesis, which cannot be fulfilled with fluorescence imaging alone. Meanwhile, in our MIP-FLF design, we relied on fluorescence intensity modulations to sense the MIP effect. As a result, we can only get MIP contrast from the molecules stained with certain fluorescent dyes, and photobleaching issue is sometimes not negligible. To mitigate these issues and broaden the potential applications, label-free single-shot volumetric chemical imaging will be pursued in our future work.

## Acknowledgments

This work is supported by R01EB032391 to J.X.C. and L.T., R35GM136223, R01AI141439, R33CA261726 to J.X.C.

