## Supplement 1 for "Single-shot Volumetric Chemical Imaging by Mid-Infrared Photothermal Fourier Light Field Microscopy"

### 1. MIP-FLF system calibration

MIP-FLF reconstruction was performed by the deconvolution of 2D measurements with depth-dependent point-spread function (PSF) of the system. Here, the PSF is displayed with the depth-dependent color map calibrated experimentally (Method). The axial range of measured PSFs determined the depth-of-focus we can restore with MIP-FLF reconstruction.

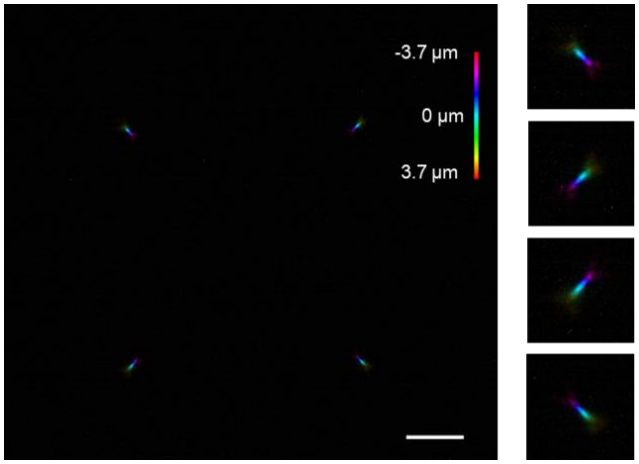


**Fig. S1.** Point-spread function (PSF) of FLF-MIP microscopy measured from 175 nm fluorescent bead, insets: zoomed in PSF.

### 2. Spatial resolution of the MIP-FLF system

The spatial resolution of MIP-FLF microscopy is determined by probe laser wavelength instead of mid-IR laser, which enables 3D mid-IR spectroscopic imaging with sub-micrometer spatial resolution. The spatial resolution of FLF microscopy is given as[1]

$$R_{xy} = \frac{\lambda N}{2NA} \qquad (S1)$$

$$R_z = \frac{d_{MLA} {R_{xy}}^2}{\lambda d_{max}} \qquad (S2)$$

where NA is the numerical aperture of MIP-FLF system, λ is the fluorescence emission wavelength (560 nm), N is the occupancy ratio (the ratio between the effective pupil size at the MLA and $d_{MLA}$), $d_{MLA}$ is the diameter of the microlens, $d_{max}$ is the distance from the outmost microlens covered by the illumination beam to the center of the microlens array (MLA). Here, NA is constrained by both the objective (0.9) and the MLA $(NA_{MLA} \times \frac{f_{MLA}}{f_{FL}} \times M = 0.97)$. M is the magnification of the objective lens, $f_{MLA}$ and $f_{FL}$ is the focal lengths of MLA and Fourier lens respectively. Consequently, the calculated lateral resolution is 580 nm and axial resolution is 850 nm according to optical propagation theory.

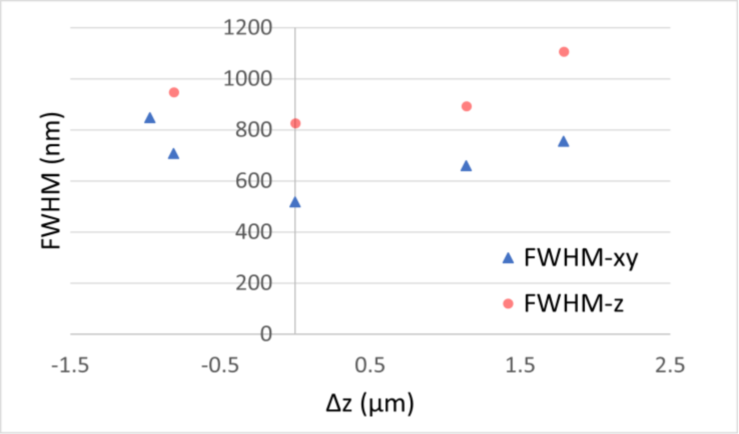


**Fig. S2.** Full width at half-maximum of the lateral and axial cross section of 175 nm fluorescent beads at varying depths.

### 3. Fourier-transform infrared spectroscopy of fatty acids

Fourier-transform infrared spectroscopy (FTIR) of $^{13}C$ labeled fatty acid mixture showed a ~30 $cm^{-1}$ peak shift to lower wavenumber compared with $^{12}C$ palmitic acid (major contents in the $^{13}C$ fatty acid mixture).

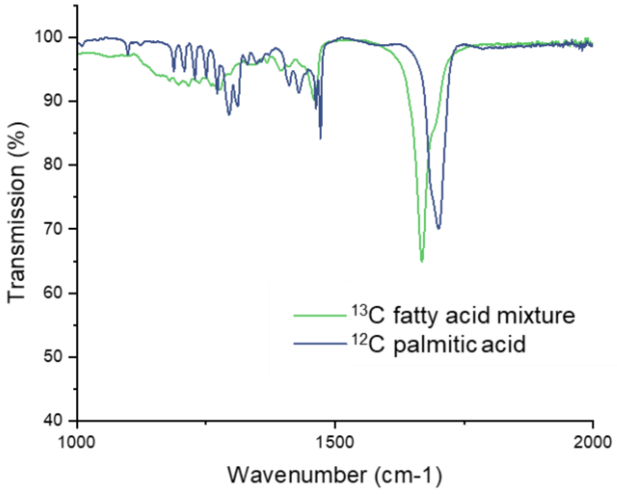


**Fig. S3.** FTIR of $^{13}C$ fatty acid mixture (green) and palmitic acid (blue).

### 4. MIP colormap of lipid contents in Mia Paca-2 and G3K cells

3D distribution of lipid contents in Mia Paca-2 cells ang G3K cells were demonstrated below. Here, green spots represent lipid droplets with MIP peak at 1744 cm$^{-1}$ ($^{12}$C=O), while red spots are lipid droplets with shifted MIP peak at 1704 cm$^{-1}$ ($^{13}$C=O). The following MIP-FLF reconstruction stacks showed the chemical mapping of additional cells in **Fig. 4c, e**.

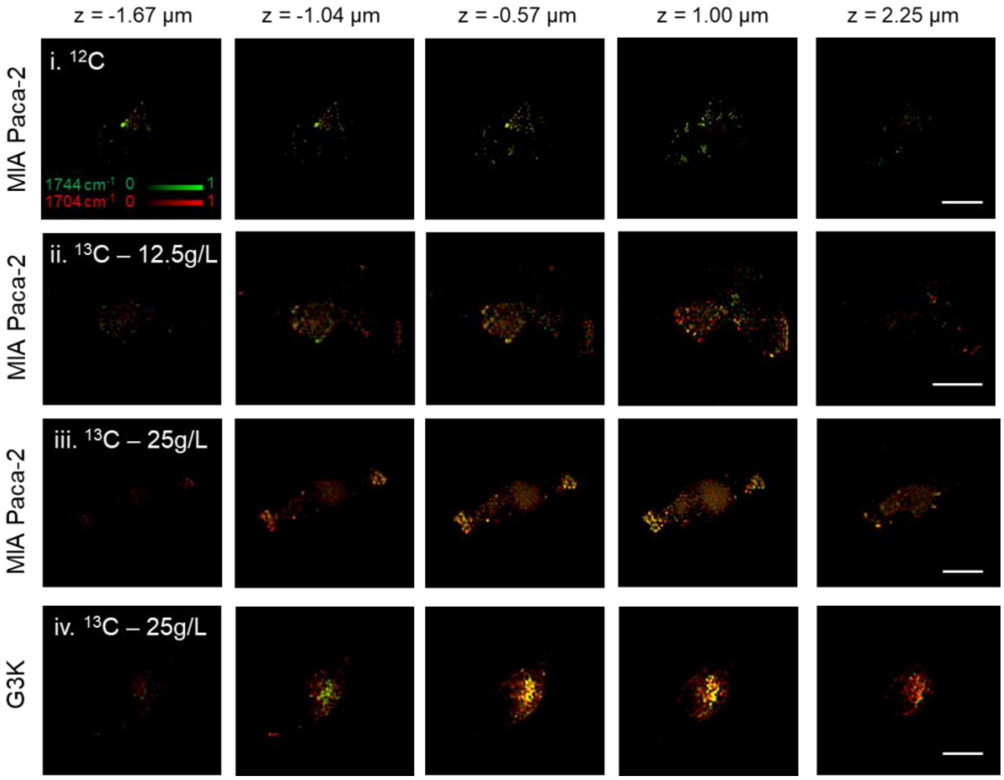


**Fig. S4.** MIP intensity colormap (red, 1704 cm$^{-1}$, green, 1744 cm$^{-1}$) at varying depths from 3D reconstructed stack of (i) MIA Paca-2 cells, (ii, iii) MIA Paca-2 cells treated with $^{13}$C fatty acids of different concentration and (iv) G3K cells treated with $^{13}$C fatty acids.
